# The Evolutionary Genomics of Grape (*Vitis vinifera* ssp. *vinifera*) Domestication

**DOI:** 10.1101/146373

**Authors:** Yongfeng Zhou, Mélanie Massonnet, Jaleal Sanjak, Dario Cantu, Brandon S. Gaut

## Abstract

We gathered genomic data from grapes (*Vitis vinifera* ssp. *vinifera*), a clonally propagated perennial crop, to address three ongoing mysteries about plant domestication. The first is the duration of domestication; archaeological evidence suggests that domestication occurs over millennia, but genetic evidence indicates it can occur rapidly. We estimated that our wild and cultivated grape samples diverged ~22,000 years ago and that the cultivated lineage experienced a steady decline in population size (*N*_*e*_) thereafter. The long decline may reflect low intensity management by humans prior to domestication. The second mystery is the identification of genes that contribute to domestication phenotypes. In cultivated grapes, we identified candidate-selected genes that function in sugar metabolism, flower development and stress responses. In contrast, candidate selected genes in the wild sample were limited to abiotic and biotic stress responses. A genomic region of high divergence corresponded to the sex determination region and included a candidate male sterility factor and additional genes with sex-specific expression. The third mystery concerns the cost of domestication. Annual crops accumulate putatively deleterious variants, in part due to strong domestication bottlenecks. The domestication of perennial crops differs from annuals in several ways, including the intensity of bottlenecks, and it is not yet clear if they accumulate deleterious variants. We found that grape accessions contained 5.2% more deleterious variants than wild individuals, and these were more often in a heterozygous state. Using forward simulations, we confirm that clonal propagation leads to the accumulation of recessive deleterious mutations but without decreasing fitness.

**Significance Statement:** We generated genomic data to estimate the population history of grapes, the most economically important horticultural crop in the world. Domesticated grapes experienced a protracted, 22,000 year population decline prior to domestication; we hypothesize that this decline reflects low intensity cultivation by humans prior to domestication. Domestication altered the mating system of grapes. The sex determination region is detectable as a region of heightened genetic divergence between wild and cultivated accessions. Based on gene expression analyses, we propose new candidate genes that alter sex determination. Finally, grapes contain more deleterious mutations in heterozygous states than their wild ancestors. The accumulation of deleterious mutations is due in part to clonal propagation, which shelters deleterious, recessive mutations.

## INTRODUCTION

The study of crop domestication has long been used as a proxy for studying evolutionary processes, such as the genetic effects of bottlenecks (1) and the detection of selection to identify agronomically important loci (2–4). Several crops have been studied in this evolutionary context (5), but there are at least two emerging issues. The first is the speed at which domestication occurs. One view, supported primarily by archaeological evidence, is that domestication is a slow process that takes millennia (6–8). Another view, based on genetic evidence and population modeling (9, 10), argues that domestication occurs much more rapidly. The gap between these two views has been bridged, in part, by a recent study of African rice. The study used population genomic data to infer that a bottleneck occurred during domestication ~3.5 thousand years ago (kya) and also that the bottleneck was preceded by a long, ~14,000 year decline in the effective population size (*N*_*e*_) of the progenitor population (11). The authors hypothesized that the protracted *N*_*e*_ decline reflects a period of low-intensity management and/or cultivation prior to modern domestication. While an intriguing hypothesis, it is not yet clear whether other crops also have demographic histories marked by protracted *N*_*e*_ declines.

The second emerging issue is the ‘cost of domestication’ (12), which refers to an increased genetic load within cultivars. This cost originates partly from the fact that the decreased *N*_e_ during a domestication bottleneck reduces the efficacy of genome-wide selection (13), which may in turn increase the frequency and number of slightly deleterious variants (14, 15). The characterization of deleterious variants is important, because they may be fitting targets for crop improvement (16). Consistent with a ‘cost of domestication’, annual crops are known to contain an increase in derived, putatively deleterious variants relative to their wild progenitors (17–20). However, it is not yet clear whether these deleterious variants increase genetic load and whether this phenomenon applies to perennial crops.

The distinction between annual and perennial crops is crucial, because perennial domestication is expected to differ from annual domestication in at least three aspects (21, 22). The first is clonal propagation; many perennials are propagated clonally but most annuals are not. Clonal propagation maintains genetic diversity in desirous combinations but also limits opportunities for sexual recombination (20, 22). The second is time. Long-lived perennials have extended juvenile stages. As a result, the number of sexual generations is much reduced for perennials relative to annual crops, even for perennials that were domesticated relatively early in human agricultural history. The third is the severity of the domestication bottleneck. A meta-analysis has documented that perennial crops retain 95% of neutral variation from their progenitors, on average, while annuals retain an average of 60% (22). This observation suggests that many (and perhaps most) perennial crops have not experienced severe domestication bottlenecks; as a consequence, their domestication may not come with a cost.

Here we study the domestication history of the grapevine (*Vitis vinifera* ssp. *vinifera*), which is the most economically important horticultural crop in the world (23). Grapes (hereafter *vinifera*) have been a source of food and wine since their hypothesized domestication ~8.0kya from their wild progenitor, *V. vinifera* ssp. *sylvestris* (hereafter *sylvestris*) (24). The exact location of domestication remains uncertain, but most lines of evidence point to a primary domestication event in the Near-East (23, 24). Domestication caused morphological shifts that include larger berry and bunch sizes, higher sugar content, altered seed morphology, and a shift from dioecy to a hermaphroditic mating system (25). There is interest in identifying the genes that contribute to these morphological shifts. For example, several papers have attempted to identify the gene(s) that are responsible for the shift to hermaphroditism, which were mapped to a ~150kb region on chromosome 2 (26, 27).

Historically, genetic diversity among *V. vinifera* varieties has been studied with simple sequence repeats (SSRs) (28). More recently, a group genotyped 950 *vinifera* and 59 *sylvestris* accessions with a chip containing 9,000 SNPs (23). Their data suggest that grape domestication led to a mild reduction of genetic diversity, indicating that grape is a reasonable perennial model for studying the accumulation of deleterious variation in the absence of a pronounced bottleneck. Still more recent studies have used whole-genome sequencing (WGS) to assess structural variation among grape varieties (29–31). Surprisingly, however, WGS data have not been used to investigate the population genomics of grapes. Here we perform WGS on a sample of *vinifera* cultivars and putatively wild *sylvestris* accessions to focus on three sets of questions. First, what do the data reveal about the demographic history of cultivated grapes, specifically the timing and severity of a domestication bottleneck? Second, what genes bear the signature of selection in *vinifera*, and do they provide insights into the agronomic shifts associated with domestication? Finally, do domesticated grapes have more derived, putatively deleterious variants relative to *sylvestris*, or have the unique features of perennial domestication permitted an escape from this potential cost?

## RESULTS

### Plant samples and population structure

We collected WGS data from nine putatively wild *sylvestris* individuals from the Near-East that represent a single genetic group (23), 18 *vinifera* individuals representing 14 cultivars, and one outgroup (*Vitis rotundifolia*) (Table S1). Our *sylvestris* accessions are a subset of the wild sample from reference (23), which was filtered for provenance and authenticity. We nonetheless label the *sylvestris* sample as ‘putatively’ wild, because it can be difficult to identify truly wild individuals. Reads were mapped to the Pinot Noir reference genome PN40024 (32), resulting in the identification of 3,963,172 and 3,732,107 SNPs across the *sylvestris* and *vinifera* samples (see Methods).

To investigate population structure, we applied principal component analysis (PCA) to genotype likelihoods (33). Only the first two principal components (PCs) were significant (*P* < 0.001); they explained 23.03% and 21.88% of the total genetic variance, respectively (Fig. 1A). PC1 separated samples of wine and table grapes, except for two accessions (Italia and Muscat of Alexandria) positioned between the two groups. PC2 divided wild and cultivated samples. Wine, table and wild grapes clustered separately in a neighbor joining tree, except for Muscat of Alexandria, which has been used historically for both wine and table grapes (Fig. 1B). Finally, STRUCTURE analyses revealed an optimal grouping of *K* = 4 which separated *sylvestris* accessions, table grapes, wine grapes and the Zinfandel/Primitivo subgroup of wine grapes, while also identifying admixed individuals (Fig. S1).

**Fig. 1.**
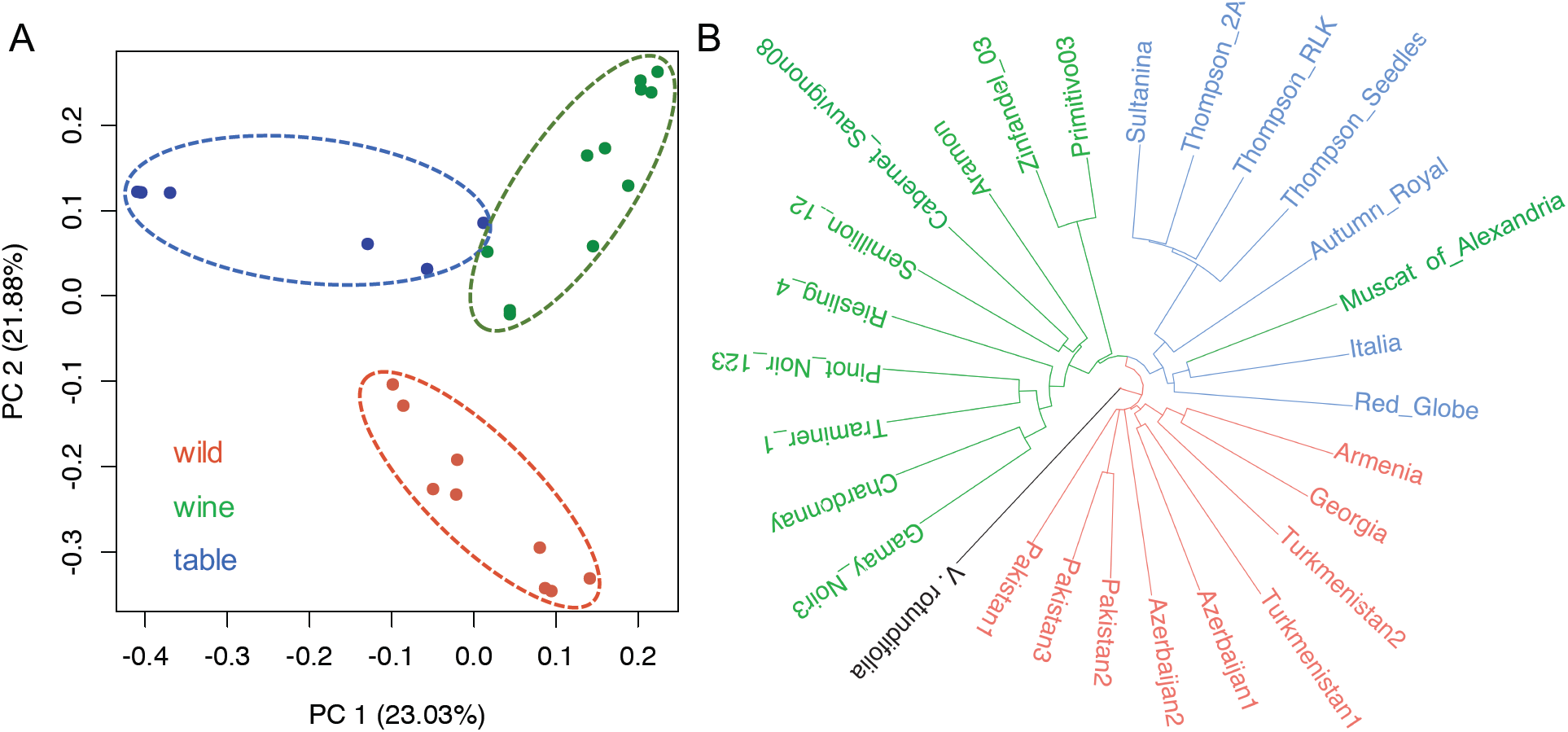
Population structure of cultivated and wild samples. (A) PCA plot based on genetic covariance among all individuals of wild samples (red) and cultivars (green for wine grapes and blue for table grapes). (B) Neighbor-Joining (NJ) tree across all samples, rooted by *V. rotundifolia.*

### Nucleotide diversity and demographic history

We estimated population genetic parameters based on the *sylvestris* accessions (*n*=9) and on a cultivated sample of *n*=14 that included only one Thompson clone and one Zinfandel/Primitivo clone (Table S1). Both samples harbored substantial levels of nucleotide diversity across all sites (*sylvestris*: π *_w_* = 0.0147 ± 0.0011; *vinifera*: π*_c_* = 0.0139 ± 0.0014; Fig. S2). Although *π* was higher in *sylvestris* (π_c_/π_w_ = 0.94 ± 0.14), *vinifera* had higher levels of heterozygosity and Tajima’s *D* values (*vinifera, D*=0.5421 ± 0.0932; *sylvestris*, *D* = -0.4651 ± 0.1577; Fig. S2). Linkage disequilibrium (LD) decayed to *r*^2^ < 0.2 within 20 kilobases (kb) in both samples, but it declined more slowly for *vinifera* after ~20kb (Fig. S2).

We inferred the demographic history of the *vinifera* sample using MSMC, which requires phased SNPs (34). Assuming a generation time of 3 years (24) and a mutation rate of 2.5×10^−9^; mutations per nucleotide per year (35), we converted scaled population parameters into years and individuals (*N*_e_). Based on these analyses, *vinifera* experienced a continual reduction of *N*_e_ starting ~22.0kya until its nadir from ~7.0kya to 11.0kya (Fig. 2A), which corresponds to the time of domestication and implies a mild domestication bottleneck. Notably, there was no evidence for a dramatic expansion of *N*_e_ since domestication. MSMC results were similar across two separate analyses (Fig. 2A), based on *n*=4 samples of either table or wine grapes (Table S1), suggesting that analyses captured shared aspects of the samples’ histories. We also used MSMC to compute divergence times. The divergence between *sylvestris* and *vinifera* was estimated to be ~22kya (Fig. 2B), which corresponds to the onset of the decline of *vinifera N*_e_. Divergence between wine and table grapes was estimated to be ~2.5kya, which is well within the hypothesized period of *vinifera* domestication (Fig. 2B).

**Fig. 2.**
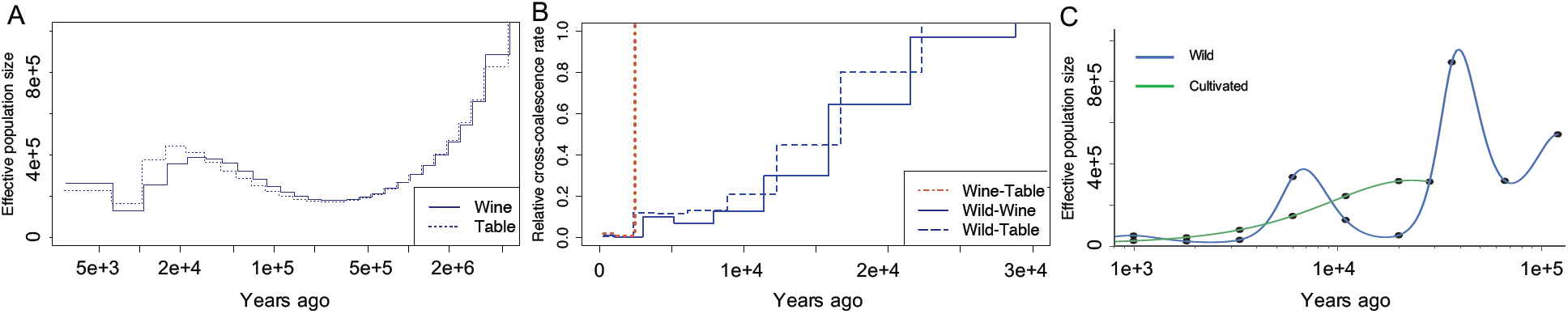
Intra- and intertaxon analyses of demographic history and divergence. (A) MSMC estimates of the effective population size (*N*_e_) *vinifera* based on two separate runs of four individuals. The solid line represents wine grapes and the dashed line is table grapes. (B) MSMC analysis of cross-coalescence (Y axis) based on comparisons between *sylvestris* and either wine or table grapes. (C) Divergence time and past *N*_e_ changes inferred by the SMC ++ analyses, based on unphased genotypes.

We repeated demographic analyses with SMC++, which estimates population histories and divergence without phasing (36)(Fig. 2C). This method yielded no evidence for a discrete bottleneck from ~7.0 to 11.0kya, but SMC++ and MSMC analyses had four similarities: *i*) an estimated divergence time (~30kya) that greatly predates domestication; *ii*) a slow decline in *vinifera N*_e_ since divergence; *iii*) no evidence for a rapid expansion in *N*_*e*_ after domestication; and *iv*) a ~2.6kya divergence of wine and table grapes (Figs. 2C & S3). We also used SMC++ to infer the demographic history of our *sylvestris* sample, revealing a complex *N*_e_ pattern that corresponds to features of climatic history (see Discussion).

### Sweep mapping

We investigated patterns of selection and interspecific differentiation across the grape genome. All sweep analyses focused on sliding 20kb windows, reflecting the genome-wide pattern of LD decline (Fig. S2). Windows that scored in the top 0.5% were considered candidate sweep regions.

We began with CLR (37) and XP-CLR (38) analyses. CLR identifies potentially selected regions by detecting skews in the site frequency spectrum (sfs) within a single taxon, while XPCLR detects sfs skews relative to a reference taxon (*sylvestris*). Within *vinifera*, CLR identified 117 20kb windows encompassing 309 candidate-selected genes (Table S2). Among those detected by CLR, nine functional categories were identified as significantly overrepresented (*P* ≤ 0.01), including the “Alcohol dehydrogenase superfamily", “Monoterpenoid indole alkaloid biosynthesis” and “Flower development” (Table S3). XP-CLR identified a similar number of genes (367); both tests identified genes involved in berry development and/or quality, including the *SWEET1* gene (Fig. S4), which encodes a bidirectional sugar transporter (39). *SWEET1* was overexpressed in full-ripe berries compared to immature berries (adj. *P*=9.4E-3; Fig. S6), suggesting an involvement in sugar accumulation during berry ripening. Additional genes of interest detected by both tests included: *i*) a leucoanthocyanidin dioxygenase (*LDOX*) gene (*VIT_08s0105g00380*) that peaks in expression at the end of veraison (adj. *P*=8.9E-10; Fig. S6) and may be involved in proanthocyanidins accumulation (40–42); *ii*) genes potentially involved in berry softening, such as two pectinesterase-coding genes and a xyloglucan endotransglucosylase/hydrolase gene that exhibited maximal expression in post-veraison berry pericarps (Fig. S6); and *iii*) flowering time genes, including a *Phytochrome C* homolog.

As a comparison, we applied CLR analyses to the *sylvestris* sample, which were notable for three reasons. First, the top 0.5% of windows yielded far fewer (88 vs. 309) genes (Table S2). Second, CLR candidate-selected regions within *sylvestris* were distinct from those in *vinifera* (Fig. 3A); none of the putatively selected regions overlapped between taxa. Third, candidate-selected genes were enriched primarily for stress resistance (Table S4), including flavonoid production (*P*=6.27E-3), ethylene-mediated signaling pathways (*P*=8.76E-6), and the stilbenoid biosynthesis pathway (*P*=1.93E-50). Stilbenoids accumulate in response to biotic and abiotic stresses (43).

**Fig. 3.**
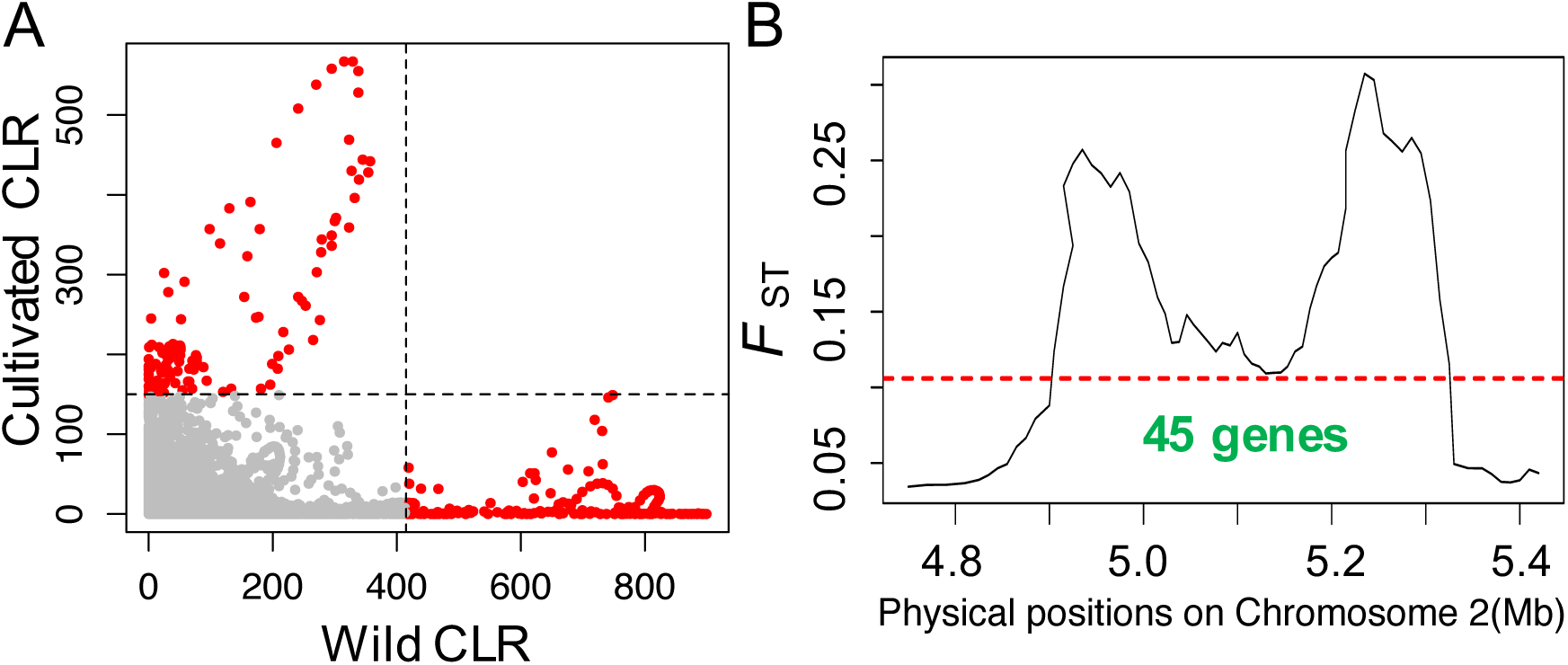
The CLR statistic computed for 20 Kbp windows along chromosomes separately for wild samples and cultivars. (A) The scatterplot shows the CLR statistic for corresponding windows for both wild (X-axis) and cultivated samples (Y-axis). The dashed line represents the 99.5% cutoff, and red dots represent outlier regions. (B) *F*_ST_ analyses between *vinifera* and *sylvestris* identify two peaks in the sex determination region that encompass 45 annotated genes.

We also detected regions of high divergence between wild and cultivated samples using *F*_*ST*_ and *D*_xy_ which identified 929 and 546 candidate-selected genes (Tables S2 & S6). A prominent region of divergence was identified by both methods from ~4.90Mb tô5.33Mb on chromosome 2 (Figs. 3B & S5), which coincides with the sex determination region (44). With both methods, the region contained two peaks of divergence. In *F*_ST_ analyses, the two peaks contain 13 and 32 genes, respectively. In the first peak, six genes were overexpressed in female (F) compared to both male (M) and hermaphroditic (H) flowers (adj. *P* ≤ 0.05; Fig. S7; Table S6), representing a non-random enrichment of F expression under the peak (binomial; *P* < 10^−7^). One of these genes had been identified as a candidate male sterility gene (*VviFSEX*) (45). The second peak included four genes with biased sex expression: one with higher F expression, two with higher H expression and one with higher M expression (Table S6).

### Deleterious variants

Domesticated annual crops accumulate more deleterious variants than their progenitors (17, 20, 46). To examine the potential increase in the number and frequency of deleterious variants at nonsynonymous sites between *vinifera* and *sylvestris* samples, we predicted deleterious SNPs using SIFT (47). A total of 33,653 nonsynonymous mutations were predicted to be deleterious in both samples. The number of derived deleterious variants was 5.2% higher, on average, for *vinifera* individuals than *sylvestris* individuals (Fig. 4), and the ratio of deleterious to synonymous variants was also elevated in *vinifera* (Fig. S8). Most (~77%) deleterious variants were found in a heterozygous state in both samples, but the distribution by state differed between taxa, because deleterious variants were more often homozygous in *sylvestris* (*P* < 0.001, Fig. 4). Cultivated accessions had a higher proportion of heterozygous deleterious variants (*P* = 0.002, Fig. 4) and an elevated ratio of deleterious to synonymous variants (*P* < 0.001, Fig. S8).

**Fig. 4.**
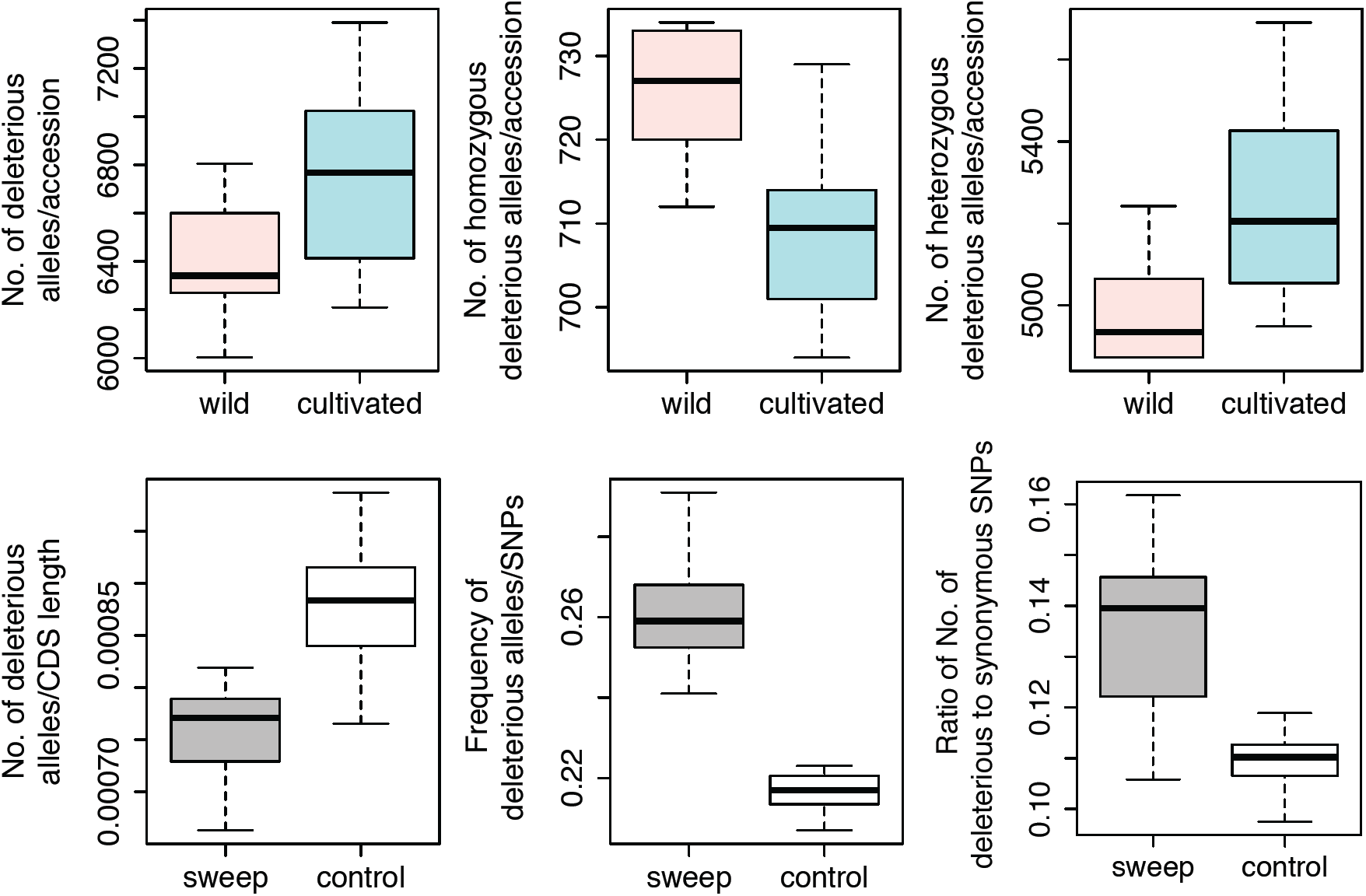
The number and frequency of derived deleterious alleles in cultivars and wild samples. Top) Comparisons between *vinifera* and *sylvestris* for the number of deleterious variants per individual overall (left), as homozygotes (middle) and as heterozygotes (right). Bottom) Comparisons between sweep regions and the rest of the genome (control) for the number (left), population frequency (middle) and ratio (right) of the number deleterious to synonymous variants per *vinifera* individual.

We also examined the distribution of putatively deleterious variants for *vinifera* in sweep regions compared to the remainder of the genome (i.e., the ‘control’). Sweep regions contained a significantly lower number of deleterious mutations when corrected for length (*P* < 0.001, Fig. 4), but these variants were also found at significantly higher frequencies (*P* < 0.001, Fig. 4) and in higher numbers relative to synonymous variants (*P* < 0.001; Fig. 4). All of these trends – including the number of deleterious variants per individual, the distribution by state, and the effects in sweep regions - were qualitatively similar using PROVEAN (48) to identify deleterious variants (Fig. S9).

Like grapes, cassava is clonally propagated, and it also has high levels of heterozygous deleterious variants (20). To determine whether clonal propagation can contribute to the accumulation of deleterious variants, we performed forward simulations under two mating systems: outcrossing and clonal propagation that began at the time of domestication (~8kya). Each mating system was considered under three demographic models: a constant size population, a long ~30ky population decline similar to that inferred from SMC++ analysis, and a discrete bottleneck (see Methods; Fig. S10). Under an additive model without back mutation, the discrete bottleneck increased the number of deleterious alleles under both mating systems but with little effect on load (Fig. S12). Under a recessive model, an outcrossing, bottlenecked population purged deleterious variants (Fig. 5) (49, 50), and clonal propagation increased the number of deleterious variants under all demographic scenarios (Fig. 5). Despite the increase in deleterious variants, clonal propagation decreased load under the recessive model (Fig. 5), because clonality hides deleterious, recessive variants.

**Fig. 5.**
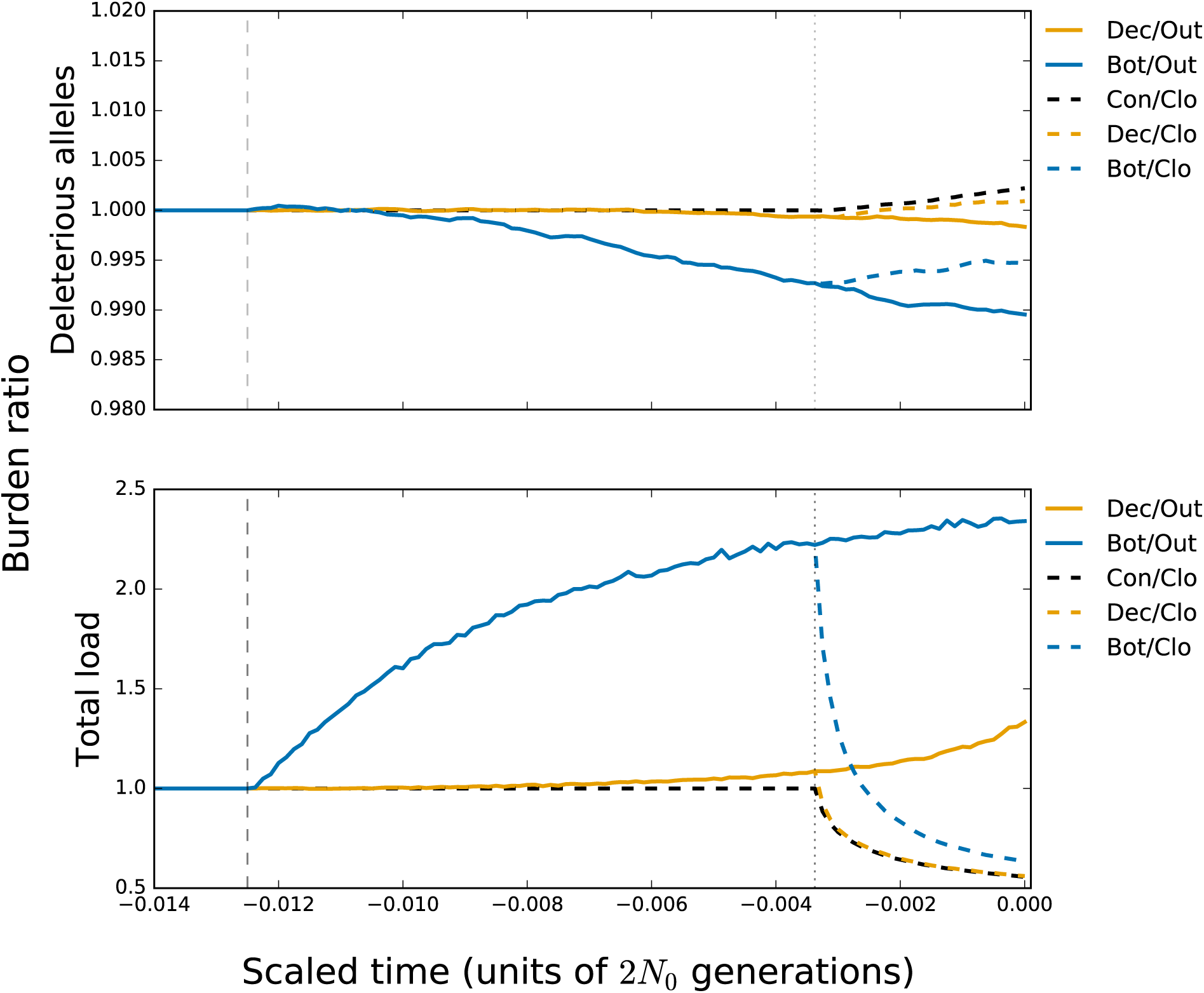
Forward simulations under a model of recessive selection for three demographic scenarios and two mating systems. The top graph represents the average number of deleterious alleles per accession relative to an outcrossing population of constant size. The bottom graph represents the total load relative to an outcrossing population of constant size. The dashed lines represent the time of demographic shift, ~30k years ago, and the onset of clonal propagation during domestication ~8k years ago. Abbreviations for demographic models are: Con = constant population size; Dec = declining population size; Bot = bottleneck. Abbreviations for mating schemes are: Out = outcrossing; Clo = clonal propagation.

## DISCUSSION

The Eurasian wild grape (*Vitis vinifera* subsp. *sylvestris*) is a dioecious, perennial, forest vine that was widely distributed in the Near East and the northern Mediterranean prior to its domestication (51). The earliest archaeological evidence of wine production suggests that domestication took place in the Southern Caucasus between the Caspian and Black Seas ~6.0-8.0kya (24, 52). After domestication, the cultivars spread south by 5.0kya to the western side of the Fertile Crescent, the Jordan Valley, and Egypt, and finally reached Western Europe by ~2.8kya (24, 53). Here, however, we are not concerned with the spread of modern grapes, but rather demographic history before and during domestication, the identity of genes that may have played a role in domestication, and the potential effects of domestication and breeding on the accumulation of deleterious variants.

### A protracted pre-domestication history?

We have gathered genome-wide resequencing data from a sample of table grapes, wine grapes and putatively wild grapes to investigate population structure and demographic history. These analyses lead to our first conclusion, which is that our *sylvestris* sample represents *bona fide* wild grapes, as opposed to feral escapees. This conclusion is evident from the fact that the *sylvestris* accessions cluster together in population structure analyses (Fig. 1), that they are estimated to have diverged from cultivated grapes ~22kya to 30kya (Fig. 2), and that the set of putatively selected genes differ markedly between the *vinifera* and *sylvestris* samples (Fig. 3A). The divergence time between wild and cultivated samples suggests, however, that our *sylvestris* accessions likely do not represent the progenitor population of domesticated grapes.

Analyses of *vinifera* data suggest that its historical *N*_*e*_ has experienced a long decline starting from ~22.0 to ~30.0kya. MSMC analyses indicate that this decline culminated in a weak bottleneck around the estimated time of domestication (Fig. 2A). The potential bottleneck corresponds to the estimated time of grape domestication and the shift from hunter–gatherer to agrarian societies (6). We note, however, that SMC++ analysis found no evidence for a distinct bottleneck, but instead inferred a consistent *N*_*e*_ decline (Fig. 2C). The question becomes, then, whether the domestication of *vinifera* included a discrete bottleneck. The evidence is mixed. The positive Tajima’s *D* for *vinifera* superficially suggests a population bottleneck, but forward simulations show that positive *D* values also result from a long population decline (Fig. S11). If there was a discrete bottleneck for grapes, we join previous studies in concluding that it was weak (23, 54, 55), based on two lines of evidence. First, the diversity level in our *vinifera* sample is 94% that of *sylvestris*, representing a far higher cultivated-to-wild ratio than that of maize (83%) (4), *indica* rice (64%) (17), soybean (83%) (56), cassava (71%) (20) and tomato (54%) (57). Second, MSMC analyses suggest a ~2 to 3-fold reduction in *N*_*e*_ at the time of domestication (Fig. 2A). This implies that 33%-50% of the progenitor population was retained during domestication, a percentage that contrasts markedly with the <10% estimated for maize (3, 58) and ~2% for rice (59).

The protracted decline in *N*_e_ for *vinifera* prompts a question about its cause(s). One possibility is that it reflects natural processes that acted on *vinifera* progenitor populations. For example, climatic shifts may have contributed to the long *N*_e_ decline, because the Last Glacial Maximum (LGM) occurred between 33.0 and 26.5kya (60). If the LGM caused *vinifera*’s population decline, one might expect to see population recovery during glacial retraction from 19.0 to 20.0kya. We detect evidence of recovery in *sylvestris* but not *vinifera* (Fig. 2C). A second possibility is that the domesticated germplasm is derived from a single deme of a larger metapopulation, because population structure can produce a signal of apparent *N*_e_ decline (61). Finally, it is possible that proto-*vinifera* populations experienced a long period of human-mediated management, as suggested in the study of African rice (11). It is difficult to prove this proposition, but three factors are consistent with this possibility: *i*) the contrasting historical pattern of the wild sample, *ii*) the fact that some sites in the Southern Caucasus mountains have evidence of human habitation for > 20k years (62) and *iii*) a growing consensus that humans altered ecosystems long before the onset of agriculture (63).

A surprising feature of demographic inference is the lack of evidence for a post-domestication expansion of *vinifera* (Fig. 2). This observation contrasts sharply with studies of maize (58) and African rice (11), both of which had > 5-fold *N*_*e*_ increases following domestication. We hypothesize that the lack of expansion in grapes relates to the dynamics of perennial domestication, specifically clonal propagation and the short time frame (in generations). Data from peach are consistent with our hypothesis, but peach also has extremely low historical levels of *N*_e_ (64). Almond, which is another clonally-propagated perennial, exhibits ~2-fold *N*_*e*_ expansion after domestication (64), but it also may have been propagated sexually prior to the discovery of grafting (65). Clearly more work needs to be done to compare demographic histories across crops with varied demographic and life histories.

Our demographic inferences have caveats. First, our study – along with all previous studies - has likely not measured genetic diversity from the precise progenitor population to *vinifera*. Indeed, such a population may be extinct or at least substantially modified since domestication. Second, our sample size is modest, but it is sufficient to infer broad historical patterns (34). Consistent with this supposition, the two runs of MSMC with two different samples of *n*=4 yielded qualitatively identical inferences about the demographic history of *vinifera*. Larger samples will be necessary for investigating more recent population history and may provide further insights into the potential for population expansion after domestication. Finally, demographic calculations assume a mutation rate and a generation time that may be incorrect, and they also treat all sites equivalently. Note, however, that masking selected regions provide similar inferences (Fig. S3) and also that our observations are consistent with independent estimates about domestication times and glacial events.

### Selective sweeps and agronomically important genes

Selective sweep analyses identified genes and regions that have been previously suspected to mediate agronomic change. One example is that of the *SWEET1* gene, which is within a potential *vinifera* sweep region. The same gene is also within a region of differentiation between non-admixed table and wine grapes (Fig. S4). Based on haplotype structures, we hypothesize that at least one difference between wine and table grapes is attributable to the *SWEET1* sugar transporter.

A major change during grape domestication was the switch from dioecy to hermaphroditism (66). The sex-determining region resides on chromosome 2, based on QTL analyses that fine-mapped the sex locus between ~4.90 and 5.05 Mbp (26, 27). The region corresponds to a larger chromosomal segment from 4.75 Mb to 5.39 Mb based on GBS data and on segregation patterns from multiple families (44). With WGS data, we have identified a similar region that contains two discrete divergence peaks, from ~4.90 to 5.05Mb and from ~5.2 to 5.3 Mb (Figs. 3B & S5). We posit that the two peaks are meaningful, because a shift to hermaphroditism may require two closely-linked loci: one that causes loss of M function and another that houses a dominant F sterility mutation (67, 68). The first peak contains six genes overexpressed in F flowers, including *VviFSEX*, which may abort stamen development (45). We predict that the second peak houses a dominant F sterility factor. The leading candidates are four genes that are differentially expressed among sexes (Table S2), but none of the four are annotated with an obvious function in sex determination (69) (Table S6).

### Putatively deleterious mutations in a clonally propagated perennial

Like grapes, most perennials have experienced moderate bottlenecks (22), raising the question as to whether they typically have an increased burden of slightly deleterious mutations (21). We find that each *vinifera* accession contains 5.2% more putatively deleterious SNPs, on average, than the wild individuals in our sample. This difference exceeds that observed for dogs (2.6%) (46) and rice (~3-4%) (17) but pales in comparison to cassava (26%), a clonally propagated annual (20). Our simulations show that clonal propagation can lead to the accumulation of deleterious recessive mutations and a reduction of load under a recessive model (Fig. 5). We do not know the dominance of variants in grapes, but we predict that most heterozygous, putatively deleterious mutations are recessive and hence do not contribute to increased load or to a cost associated with domestication. These same mutations do, however, provide a genomic explanation a well-known feature of grape breeding: severe inbreeding depression (70).

## MATERIALS AND METHODS

For full materials and methods, please see SI Appendix, Supplementary Text. We collected leaf tissue for 13 individuals from 11 *vinifera* cultivars, nine *sylvestris* accessions and one accession of *V. rotundifolia* (Table S1). DNA was extracted from leaf samples, Illumina paired-end sequencing libraries were constructed (TrueSeq), and libraries were sequenced as 150-bp paired reads. Illumina raw reads for five other cultivars were gathered from the Short Read Archive at NCBI (Table S1).

Reads were trimmed, filtered and mapped to the PN40024 reference (12X) (32). Local realignment was performed around indels; reads were filtered for PCR duplicates; and sites with extremely low or high coverage were removed. For population structure analyses, we used ANGSD (33) to generate a BEAGLE file for the variable subset of the genome, and then applied NGSadmix (71). To measure genome-wide genetic diversity and other population parameters, we estimated a genome-wide sfs from genotype likelihoods (33).

Functional regions were based on the *V. vinifera* genome annotation in Ensembl (v34). Nonsynonymous SNPs were predicted to be deleterious based on a SIFT score ≤ 0.05 (72). The *V. rotundifolia* outgroup allele was submitted to prediction programs to avoid reference bias (17, 18). The number of deleterious or synonymous alleles per individual or region was calculated as 2 × the number of homozygous variants + heterozygous variants (73).

We employed MSMC 2.0 to estimate *N*_e_ over time (34, 74), based on SNPs called in GATK v3.5 (75)(see SI Appendix, Supplementary Text). Segregating sites within each sample were phased and imputed using Shapeit (76) based on a genetic map (44). Demographic history was also inferred with SMC++, which analyzes multiple genotypes without phasing (36). SweeD (37) and XP-CLR (38) were used to detect selective sweeps. *F*_ST_ and *D*_xy_ values were averaged within 20 kbp non-overlapping windows using ANGSD (33).

Functional categories were assigned to genes using VitisNet functional annotations (77). We tested functional category enrichment using Fisher’s Exact Test, with *P* ≤ 0.01 as significant. Gene expression data used SRA data for berry (SRP049306) and flower (SRP041212) samples. Reads were trimmed for quality and mapped onto the PN40024 transcriptome (v.V1 from http://genomes.cribi.unipd.it/grape/) using Bowtie2 (78). DESeq2 (79) was used to normalize read counts and to test for differential expression.

Forward in time simulations were carried out using fwdpy11 (80). Five hundred replicate simulations were run for each demographic and mating scheme model. The population decline model was based on SMC++ results and rescaled for computational performance. Three demographic models were simulated: constant population size, a linear population decline, and a discrete bottleneck. Two mating schemes were simulated: strict outcrossing for the whole simulation, and outcrossing at the onset of domestication. Additional details are available from SI Appendix, Supplementary Text.

## ACKNOWLEDGEMENTS

We thank two anonymous reviewers, R. Gaut and R. Figueroa-Balderas for generating the data and sampling and also D. Seymour, Q. Liu, K. Roessler and E. Solares for comments. YZ is supported by the International Postdoctoral Exchange Fellowship Program; JS is supported by the NSF-GRFP; BSG is supported by the Borchard Foundation. DC is supported by J. Lohr Vineyards and Wines and by E. & J. Gallo Winery.

